# Non-linear growth models combined with survival rates identify the optimal rearing temperature and harvest timing in edible crickets, *Gryllus bimaculatus* and *Teleogryllus occipitalis*

**DOI:** 10.64898/2026.09.16.751979

**Authors:** D. Akiyama, K. Murata, M. Akechi, Y. Morimitsu, K. Yura, H. Hirooka, T. Suzuki

## Abstract

Crickets are gaining attention as sustainable protein sources because of their high nutritional value and low environmental impact. Growth models are essential tools for optimising production by predicting growth characteristics. However, previous studies have not compared multiple non-linear growth models for crickets or explicitly considered environmental factors. We analysed the growth patterns of the two-spotted cricket (*Gryllus bimaculatus*) and the Asian cricket (*Teleogryllus occipitalis*) at multiple temperatures (25.0, 27.5, 30.0, 32.5 and 35.0 °C) using three non-linear models (Gompertz model, logistic model, and von Bertalanffy model) and combined the predicted body mass with observed survival rates to predict the optimal harvest time and temperature. All three models fitted the data well (*R*^2^ ≥ 0.97 in both species). The asymptotic body mass was highest at 30.0 °C, and the growth rate constant increased with temperature, reaching its maximum at 35.0 °C except for the logistic model in *G. bimaculatus* (32.5 °C). In addition, the predicted value of harvest, calculated by combining predicted body mass and observed survival probability, was highest at 30.0 °C in both species, reaching 13.82 kg/m^3^ at 38 days in *G. bimaculatus* and 7.12 kg/m^3^ at 62 days in *T. occipitalis*. These results suggest that 30.0 °C is the most suitable constant-temperature condition for maximising predicted productivity, although the optimal harvest timing differs between species.

## Introduction

As the global population grows, the demand for food and feed is predicted to increase (United Nations, 2022). In addition to expanding the production of current food and feed sources, various alternatives are needed to diversify supply and build a resilient food system. Insects are among the promising options (van Huis *et al*., 2013). Insect farming uses less land, water, feed, and other resources and produces fewer greenhouse gas emissions than conventional livestock farming (Dobermann *et al*., 2017). Insects are also known to be highly nutritious, containing high levels of protein, fat, minerals, vitamins and energy (Ramos-Elorduy *et al*., 1997). Currently, commercial farming of insects as food and feed targets approximately 10 different species. Among them, crickets are among the most widely farmed because they have a mild flavour suitable for a variety of dishes (Van Itterbeeck and Pelozuelo, 2022). However, mass production of crickets faces several challenges, with the lack of standardised feeding protocols being one of the most important. Because crickets are omnivorous and can consume diverse types of food, diets vary considerably across farms, resulting in a shortage of robust growth models that can reliably predict production outcomes. Most cricket farms rely on poultry or aquaculture feeds because of their easy availability and in response to empirical observations of relatively good growth performance (Hanboonsong and Durst, 2020). However, feed costs account for 50% to 60% of the total production costs, highlighting the need for improvement (Cansee *et al*., 2025). Temperature is another critical factor in farming, as it affects growth rate, yield and feed consumption (Kong *et al*., 2025; Magara *et al*., 2024). In non-tropical regions, maintaining optimal temperatures requires substantial heating, directly increasing production costs.

Growth models are essential tools for stable production and planned shipments. Using an optimal model enables the prediction of growth rates and feed consumption at various stages, supporting decision-making during production (Wong *et al*., 2025). Various non-linear models have been studied in the livestock and aquaculture industries (Sun and Wang, 2024; Teleken *et al*., 2017; Tsukahara *et al*., 2008). Furthermore, the parameters obtained from growth curves provide information related to the growth characteristics. Knowledge of factors affecting the shape of the growth curves and the relationship between the parameters is required to improve efficiency (Morrow *et al*., 1978).

In crickets, Kaewtapee *et al*. (2024) estimated the optimal protein requirement for the two-spotted cricket *Gryllus bimaculatus* by using a Gompertz curve, whereas Mauritsson and Jonsson (2023) reported a mechanism-based model for maintenance and feeding of the house cricket *Acheta domesticus*. However, previous studies have generally focused on a single model and have not conducted comparisons across multiple growth models. In addition, they have not explicitly considered the effect of temperature on cricket growth; this limits their applicability under different temperature conditions.

Growth studies of crickets have mainly used two major species, *G. bimaculatus* and *A. domesticus*. On the other hand, the Asian cricket *Teleogryllus occipitalis*, distributed in Southeast and East Asia including the southern part of Japan, is also commonly consumed in Asia (Hanboonsong *et al*., 2013). Including a wider variety of cricket species in growth model studies is valuable for comparing rearing features and ensuring the diversity of edible cricket options.

The purpose of this study was to analyse the growth patterns of two species of edible crickets under multiple temperature conditions using non-linear models. Two aspects were considered: (1) the effect of temperatures on growth curve parameters using three growth curve models; and (2) the optimal harvest time and temperature predicted by combining the growth models with observed survival rates.

## Materials and methods

### Experimental design

We used a population of *G. bimaculatus* purchased from Tsukiyono Farm Co., Ltd. (Gunma, Japan) and *T. occipitalis* collected on Amami Ohshima (Kagoshima, Japan) and previously used for whole-genome sequencing (Kataoka *et al*., 2020). The population was maintained in a wooden container (42 × 53 × 48 cm, 107 L) in which poultry feed (Chougennki Edsukeyousuuyou; Nosan Corp., Kanagawa, Japan) and a water-filled polystyrene cup (662 mL) with a paper plug were provided. We also provided the same size of cup containing water-soaked cotton wool, allowing female adult crickets to lay their eggs. Eggs laid in the water-soaked cotton wool were collected every week and kept in an incubator maintained at 30 °C until hatching. Nymphs hatched within 24 h were used for the following experiments.

In accordance with the method of Murata *et al*. (2025), 50 randomly selected nymphs were placed in a polypropylene container (239 × 176 × 91 mm; Nakaya Kagaku Sangyo, Osaka, Japan), in which the population density was approximately 15,800 crickets/m^3^, excluding the volume of the polystyrene cup used for serving water. Two pieces of pulp egg carton (100 × 150 mm) were placed in the container as shelters. A polystyrene Petri dish was placed on top of the pulp egg carton as a feeding area. The cricket feed used was based on the composition that demonstrated the highest growth performance (crude protein 22%) in the study by Kaewtapee *et al*. (2024). Details of the formulation and nutritional values are given in Table S1. Each feed ingredient was ground in a grinder (B0BRQ4NSMW; Kalelaisu, China) and sieved through a 1.0-mm sieve to minimise particle size variation. To ensure easy access to water and food, paper towelling (Nippon Paper Crecia Co., Ltd.; Tokyo, Japan) was placed in the vicinity of the feed dish and the watering device. To prevent escape, non-woven fabric (MonotaRO Co., Ltd., Osaka, Japan) was placed between the container and the lid, which had about 100 ventilation holes (2.5 mm diameter). Detailed information on the rearing setup is provided in Figure S1.

The crickets were reared in an incubator maintained at 25.0, 27.5, 30.0, 32.5 or 35.0 °C to create growth curve models. The photoperiod was maintained at light:dark = 12:12. Ten crickets were selected randomly from each container, and their body mass was recorded every week from the day of hatching (defined as day 0, hereafter d0) to the final recording day. The number of surviving crickets in each container was also counted weekly from d0 to the final recording day, and these counts were used to calculate survival probability. Hereafter, age in days after hatching is abbreviated as ’d’ followed by the number of days (e.g., d42 = 42 days after hatching). The final day of body mass recording was d70, d49, d42, d42 and d42 for *G. bimaculatus* and d140, d91, d70, d77, and d56 for *T. occipitalis* in the 25.0, 27.5, 30.0, 32.5 and 35.0 °C treatments, respectively, by which time most crickets had reached adulthood (>85% of randomly sampled individuals). The body mass data from d0 to the final recording day in each treatment were used for model fitting and predicted value of harvest (PVH) calculation (described below). The body mass data of the day of the body mass peak were used for adult body mass evaluation. This parameter was compared within the same sex between temperature treatments by statistical analyses (see ’*Statistical analyses*’ for the definition of the peak day and of significant differences). Data were collected from four independent experimental runs. Owing to a handling error, only three replicates were conducted for *G. bimaculatus* at 27.5 °C.

### Growth curve models

We employed three types of non-linear growth models: logistic, Gompertz, and von Bertalanffy. Initial parameter values were determined individually for each temperature treatment by applying a logarithmic transformation. The initial value of asymptotic body mass (*A*0) was set to 1.05 times the maximum observed body mass in each temperature treatment. With *A* fixed at this initial value (*A*0), the transformed body mass was subjected to regression analysis against age to obtain initial estimates of the remaining parameters *B* and *k* (Köksal Babacan and Demir, 2024). The calculation formulas for each model are shown in Table 1. The calculated initial values are listed in Table S2. The parameters of each model were estimated by non-linear least squares using the Gauss–Newton algorithm, implemented through the nls function in R (R Core Team, 2025).

**TABLE 1.** Growth curve models and linearised form for initial-value calculation.

| Model | Growth curve equation<br>(non-linear expression) | Linear regression equation for initial values<br>of $B$ and $k$ |
| --- | --- | --- |
| logistic | $W(t) = \left( \frac{A}{1 + Be^{-kt}} \right)$ | $\ln \left( \frac{A0}{W(t)} - 1 \right) = \ln(B) - kt$ |
| Gompertz | $W(t) = Ae^{(-B \times e^{-kt})}$ | $\ln \left[ -\ln \left( \frac{W(t)}{A0} \right) \right] = \ln(B) - kt$ |
| von Bertalanffy | $W(t) = A(1 - Be^{-kt})^3$ | $\ln \left[ 1 - \left( \frac{W(t)}{A0} \right)^{\frac{1}{3}} \right] = \ln(B) - kt$ |

$W(t)$ , body mass at a given age (day); $t$ , age (days); $A$ , asymptotic body mass (mg); $B$ , parameter that maintains the shape of the model; $k$ , growth rate constant; $A0$ , initial value of asymptotic body mass.

For the logistic and Gompertz models, the age at the inflection point was calculated by using Equation (1) (Juárez-Caratachea *et al*., 2019). For the von Bertalanffy model, the age at the inflection point was calculated using Equation (2) (Zárate-Contreras *et al*., 2022).

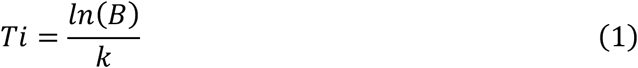

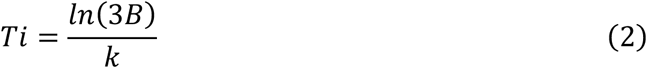

where *Ti* is the age at the inflection point, *B* is a parameter that maintains the shape of the model, and *k* is the growth rate constant.

The model’s goodness of fit was evaluated by using the coefficient of determination (*R*^2^). The *R*^2^ value is calculated by using Equation (3) and is a commonly used comprehensive indicator for measuring the accuracy of predictions (Yang *et al*., 2016). An *R*^2^ value of 1 indicates that the values predicted by the model perfectly match the actual values.

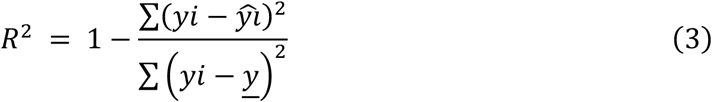

where *yi* is the observed cricket body mass, *ŷi* is the estimated cricket body mass of *yi* and *y* is the mean of the observed cricket body mass.

### Evaluation of productivity

Survival probability was determined from the observed mean number of surviving crickets at each observation age. For ages on which survival was not directly observed, survival probability was estimated by linear interpolation between two consecutive observation points. The survival probability at age t was calculated using Equation (4):

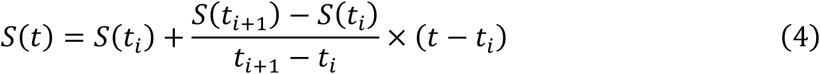

where *S* (*t*) is the survival probability at age *t*, *t*_i_ and *t*_i+1_ are two consecutive observation ages, *S* (*t_i_*) and *S* (*t_i_*_+1_) are the observed average survival probabilities at these respective time points.

The predicted value of harvest (*PVH*) was calculated by combining the predicted body mass from the logistic growth model with the survival probability obtained from Equation (4). Specifically, yield density at age *t* was estimated using Equation (5).

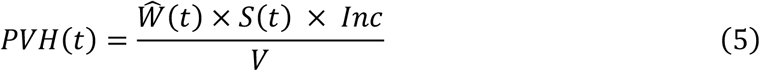

where *PVH* (*t*) is the predicted value of harvest at age *t*, *Ŵ* (*t*) is the predicted body mass obtained from the logistic growth model, *S* (*t*) is the survival probability obtained from Equation (4), *Inc* is the initial number of crickets, and *V* is the rearing container volume (m^3^). In this study, *Inc* was set to 50 and *V* to 0.003 m^3^. The volume of the container excluding the polystyrene cup used for serving water was 0.00317 m^3^, which was rounded down to 0.003 m^3^ to account for the space occupied by the egg cartons and the feed dish.

### Statistical analyses

To decide on the day of peak body mass, the means of the crickets’ body mass were statistically analysed between the previous and the current week under the same temperature treatment, by using Student’s *t*-test, Welch’s *t*-test, the Wilcoxon-Mann-Whitney *U*-test and the Brunner-Munzel test, according to the data distribution (Shapiro-Wilk test) and the homogeneity of variance (Bartlett test). To test for significant differences in adult body mass between multiple temperature treatments, the Wilcoxon-Mann-Whitney *U*-test (Benjamini-Hochberg adjusted) and the Tukey HSD test were used, according to the data distribution and the homogeneity of variance. Statistical differences in the Kaplan–Meier survival curves from d0 to the final recording day were analysed by using a pairwise log-rank test with Benjamini-Hochberg adjustment.

## Results

### Growth characteristics

The growth curves of *G. bimaculatus* and *T. occipitalis* reared at various air temperatures are shown in Figures 1 and 2, respectively. For *G. bimaculatus*, the estimated parameters for the growth curve models are shown in Table 2. Similarly, for *T. occipitalis*, the model parameters are shown in Table 3. The three growth models used in this study fitted the data well for both *G. bimaculatus* and *T. occipitalis*. For *G. bimaculatus*, the logistic model fitted best at 25.0 and 32.5 °C, the logistic and Gompertz models fitted equally well at 27.5 and 30.0 °C, and the Gompertz model fitted best at 35.0 °C (Table 2, Figure 1). The asymptotic body mass (*A*) was highest at 30.0 °C, whereas the growth rate constant (*k*) increased with temperature and was highest at 35.0 °C in the Gompertz and von Bertalanffy models and at 32.5 °C in the logistic model. For *T. occipitalis*, the logistic model fitted best at 27.5 and 30.0 °C, whereas the Gompertz model fitted best at 25.0, 32.5, and 35.0 °C (Table 3, Figure 2). A similar temperature dependence of *A* and *k* was observed in *T. occipitalis*, with *k* being highest at 35.0 °C in all three models (Table 3).

**FIGURE 1.**
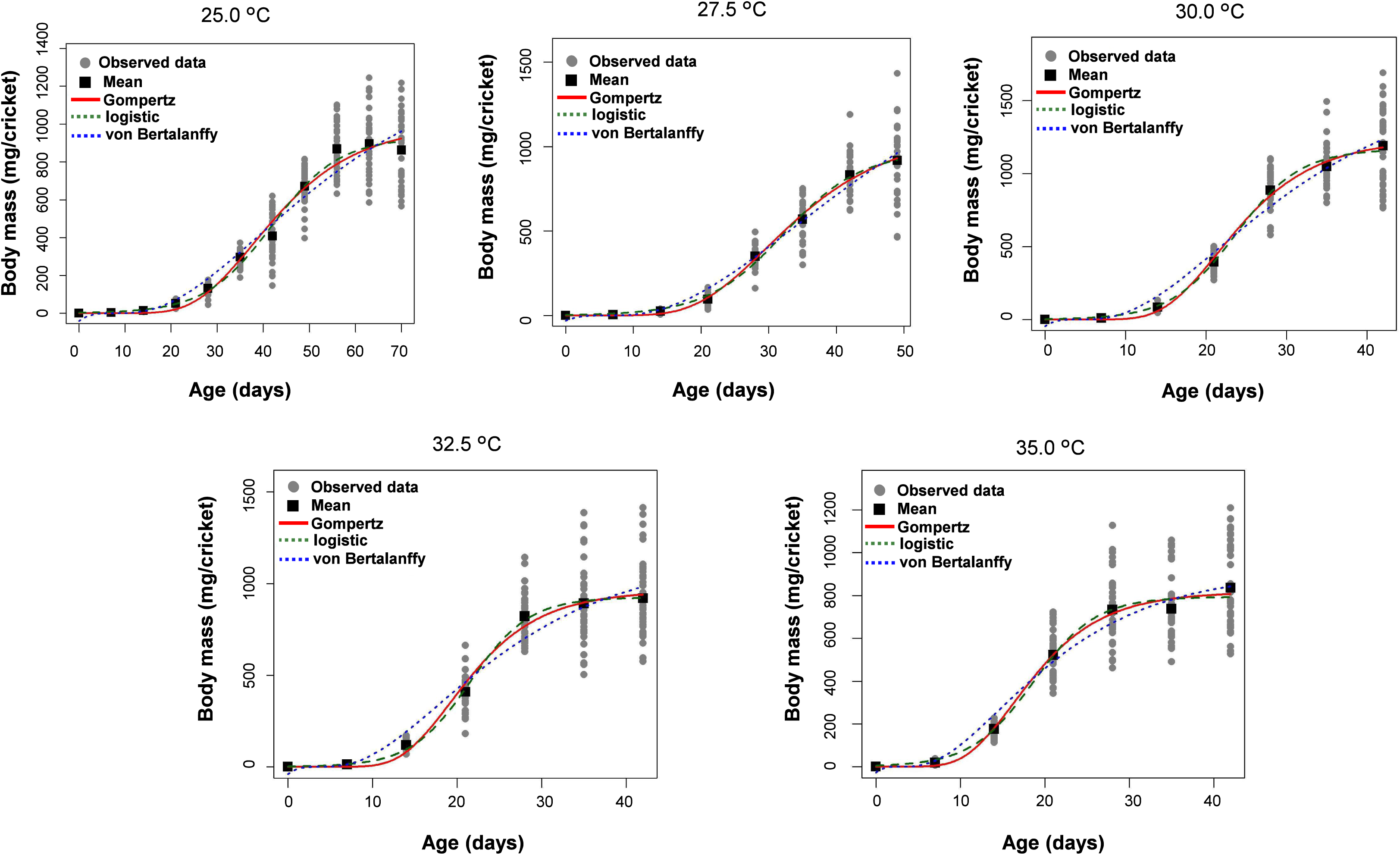
Growth curves of the two-spotted cricket, *Gryllus bimaculatus*, reared at air temperatures of 25.0, 27.5, 30.0, 32.5, and 35.0 °C. Grey circles indicate observed data and black squares indicate means. Lines show the fitted Gompertz (red solid), logistic (green dashed) and von Bertalanffy (blue dotted) models.

**FIGURE 2.**
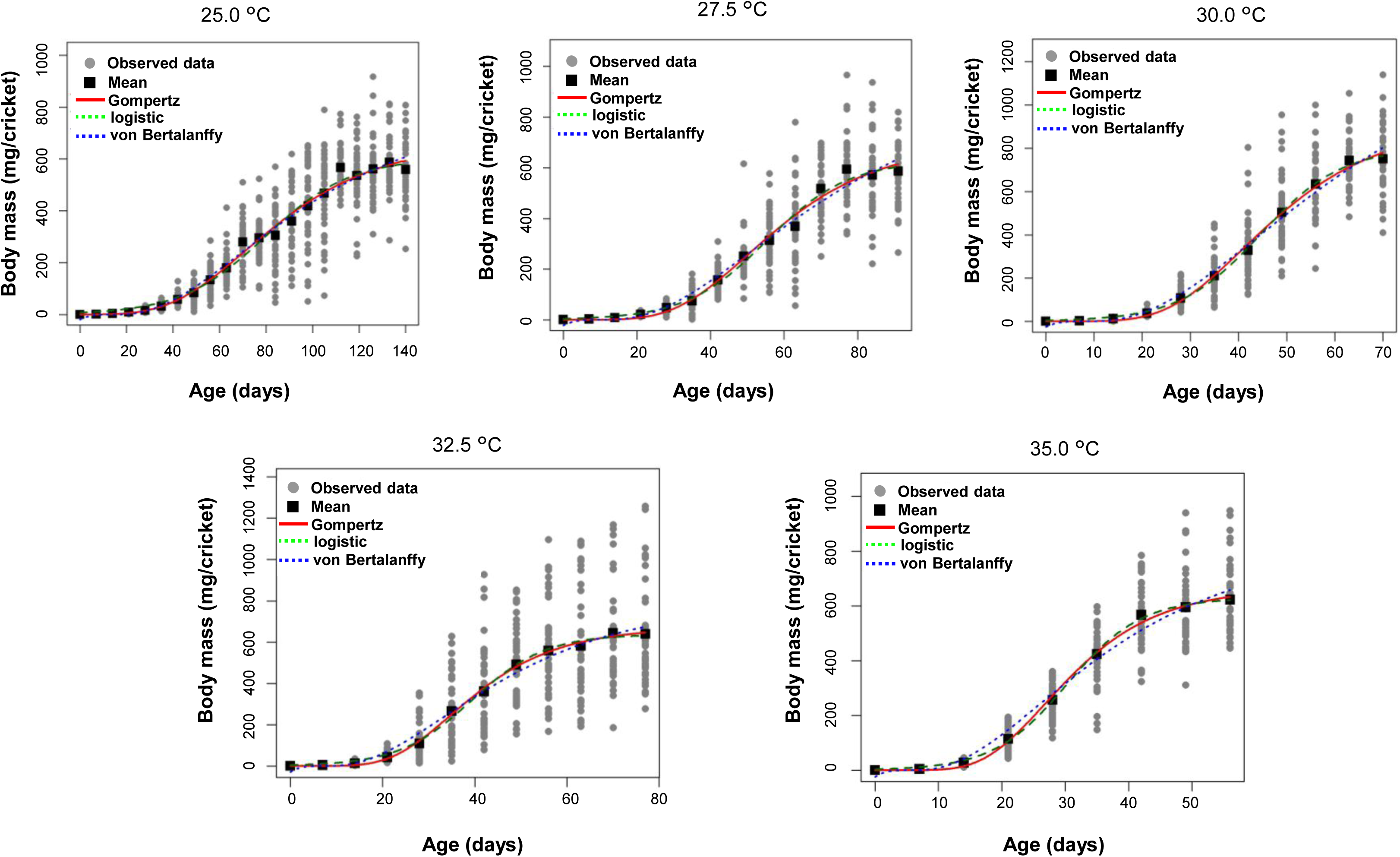
Growth curves of the Asian cricket, *Teleogryllus occipitalis*, reared at air temperatures of 25.0, 27.5, 30.0, 32.5, and 35.0 °C. Grey circles indicate observed data and black squares indicate means. Lines show the fitted Gompertz (red solid), logistic (green dashed) and von Bertalanffy (blue dotted) models.

**TABLE 2.** Parameters of *Gryllus bimaculatus* growth curve models at air temperatures of 25.0, 27.5, 30.0, 32.5, and 35.0 °C.

| | | $A$ | $B$ | $k$ | $T_i$ | $R^2$ |
| --- | --- | --- | --- | --- | --- | --- |
| 25.0 °C | logistic | 926.970 | 323.223 | 0.13793 | 41.8933 | 0.992 |
|  | Gompertz | 996.463 | 20.580 | 0.07996 | 37.823 | 0.987 |
|  | von Bertalanffy | 1371.749 | 1.311 | 0.03520 | 38.903 | 0.973 |
| 27.5 °C | logistic | 970.877 | 274.429 | 0.17404 | 32.2609 | 0.997 |
|  | Gompertz | 1084.605 | 17.877 | 0.09726 | 29.6475 | 0.997 |
|  | von Bertalanffy | 1764.776 | 1.258 | 0.03934 | 33.760 | 0.989 |
| 30.0 °C | logistic | 1165.123 | 362.893 | 0.24900 | 23.6711 | 0.997 |
|  | Gompertz | 1234.446 | 24.511 | 0.14922 | 21.439 | 0.997 |
|  | von Bertalanffy | 1649.723 | 1.300 | 0.06300 | 21.603 | 0.983 |
| 32.5 °C | logistic | 923.687 | 473.091 | 0.28621 | 21.5202 | 0.999 |
|  | Gompertz | 958.919 | 27.122 | 0.17144 | 19.2507 | 0.993 |
|  | von Bertalanffy | 1152.713 | 1.326 | 0.07718 | 17.890 | 0.973 |
| 35.0 °C | logistic | 793.444 | 156.738 | 0.27097 | 18.6536 | 0.994 |
|  | Gompertz | 815.842 | 17.397 | 0.17451 | 16.3675 | 0.995 |
|  | von Bertalanffy | 913.670 | 1.314 | 0.09296 | 14.756 | 0.984 |

$A$ is the asymptotic body mass (mg), $B$ is a parameter that maintains the shape of the model, $k$ is the growth rate constant, and $T_i$ is the age at the inflection point (days).

**TABLE 3.**
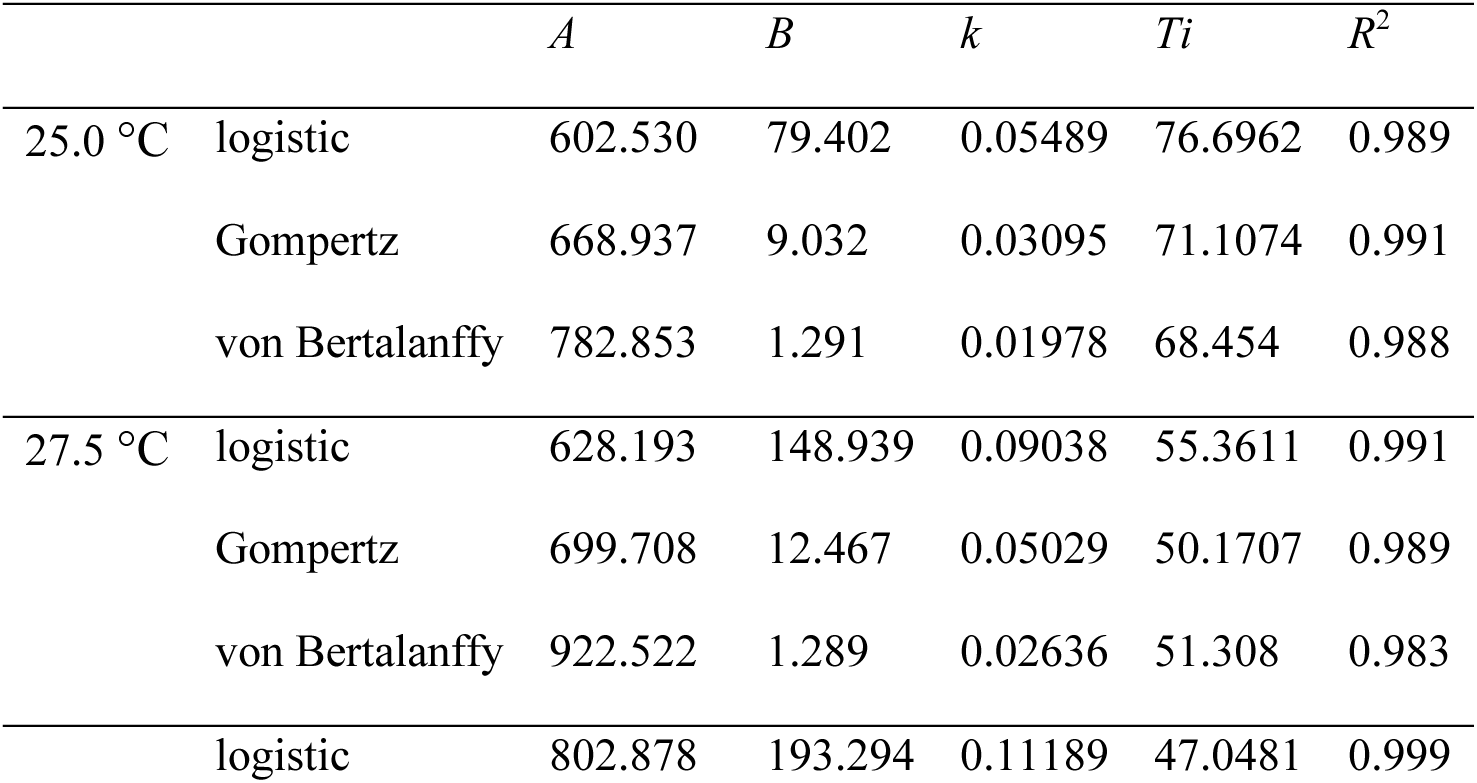
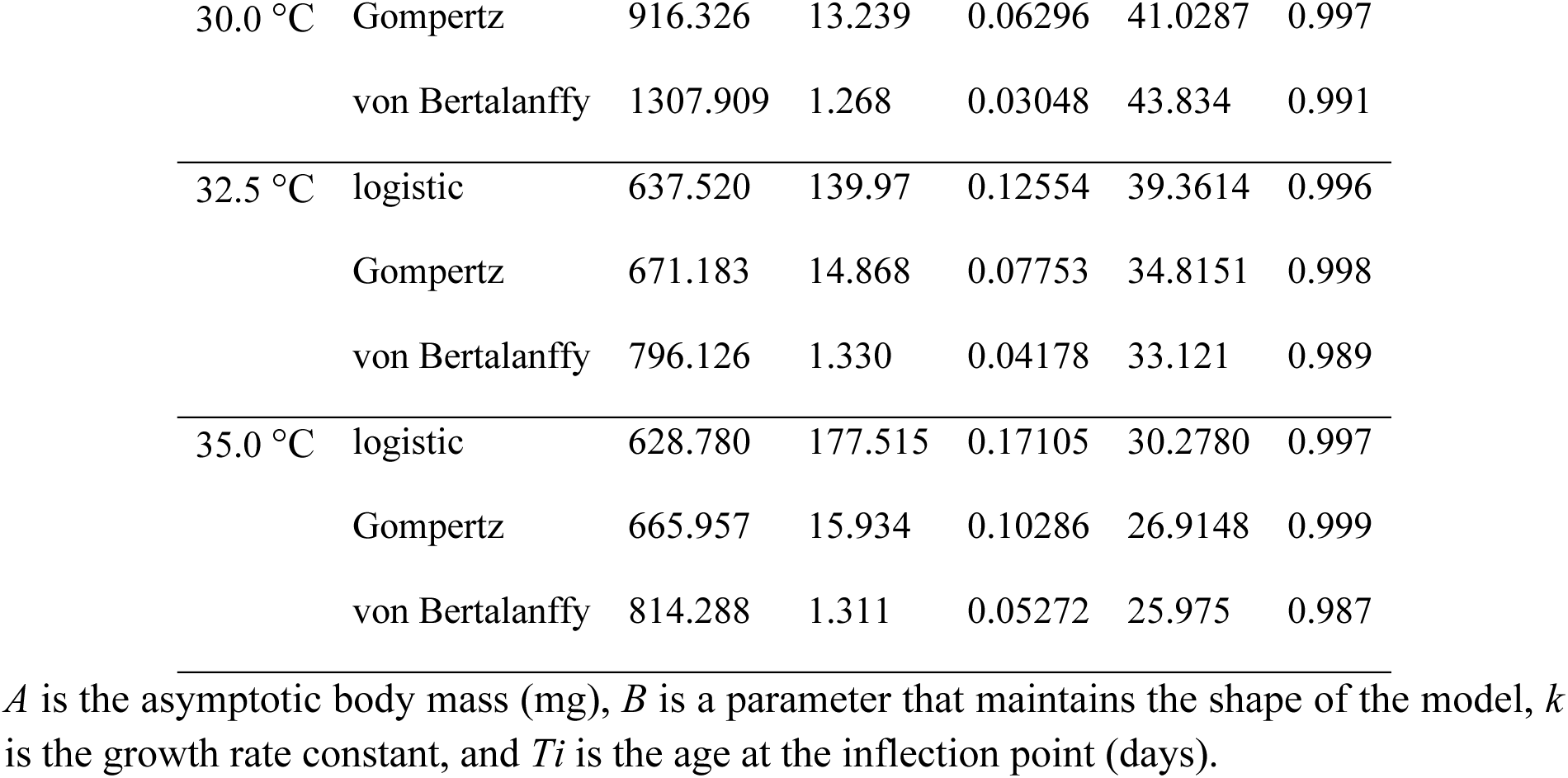
Parameters of *Teleogryllus occipitalis* growth curve models at air temperatures of 25.0, 27.5, 30.0, 32.5, and 35.0 °C.

### Body mass and survival

In *G. bimaculatus*, the first adults were observed on d56, d42, d28, d28 and d35 in the treatments of 25.0, 27.5, 30.0, 32.5, and 35.0 °C, respectively. The peak of adult body mass in the observed data (25.0 °C: d63; 27.5 °C: d49; 30.0 °C: d42, 32.5 and 35.0 °C: d35) was significantly higher in the 30.0 °C treatment (*P* < 0.05) than in the other treatments in females (Figure 3A) and than in most other treatments in males (Figure 3B). Almost all (>85%) individuals that we picked randomly for body mass measurement had reached the adult stage at the time of the peak body mass. At the time of peak body mass, females were significantly heavier than males in all temperature treatments (*P* < 0.05; 25.0 and 27.5 °C, Student’s *t*-test; 30.0, 32.5 and 35.0 °C, Welch’s *t*-test). Survival of nymphs was highest at 25.0 °C and lowest at 32.5 °C, although survival at 25.0 °C did not differ significantly from that at 30.0 °C (Figure 3C).

**FIGURE 3.**
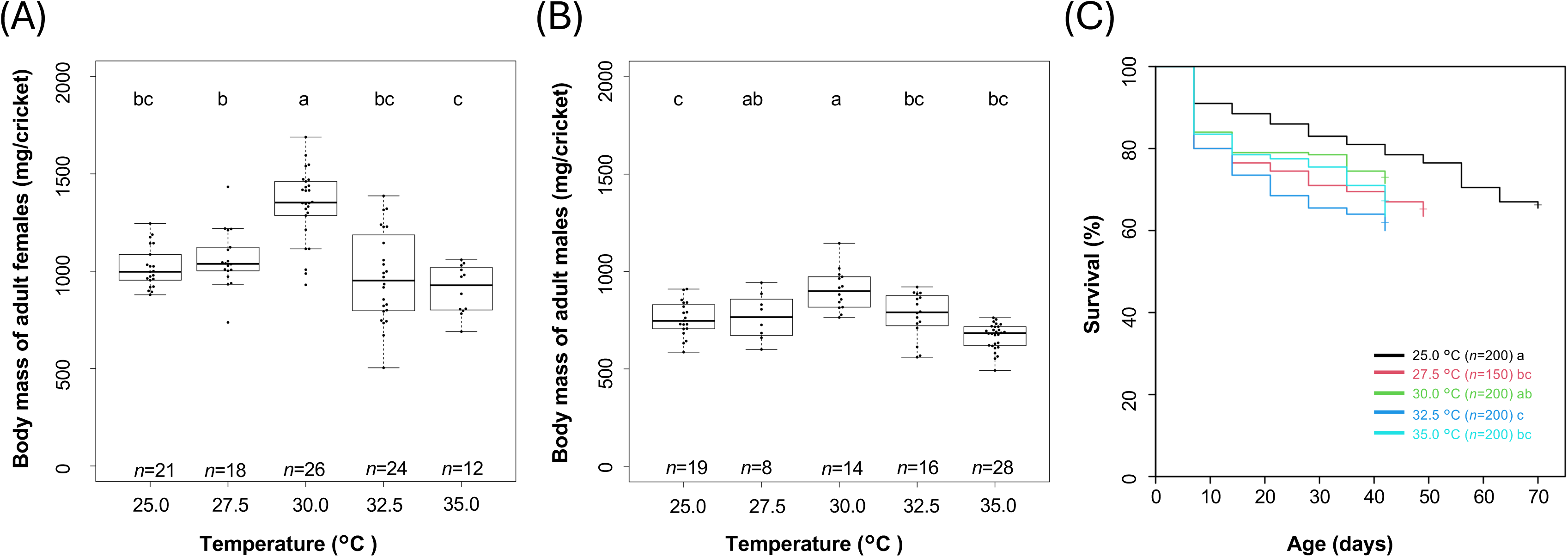
Growth performance of *Gryllus bimaculatus*. (A) Female and (B) male adult body mass at each temperature (25.0, 27.5, 30.0, 32.5, and 35.0 °C) on the day of peak body mass (d63, d49, d42, d35, and d35, respectively). Different letters indicate significant differences among temperatures (*P* < 0.05). (C) Survival rate at each temperature from d0 to d70, the final recording **day at 25.0 °C. Different letters indicate significant differences among survival curves (pairwise log-rank test, *P* < 0.05).**

In *T. occipitalis*, the first adults were observed on d77, d63, d49, d42 and d42 in the treatments of 25.0, 27.5, 30.0, 32.5, and 35.0 °C, respectively. The peak of adult body mass in the observed data (25.0 °C: d133; 27.5 °C: d91; 30.0 °C: d70, 32.5 °C: d77 and 35.0 °C: d49) tended to be higher in the 30.0 °C treatment than in the other treatments in both females (Figure 4A) and males (Figure 4B) although not all pairs between 30.0 °C and other treatments showed significant differences. Almost all (>85%) individuals that we picked randomly for body mass measurement had reached the adult stage at the time of the peak body mass. At the time of peak body mass, females were significantly heavier than males in all temperature treatments (*P* < 0.05; 27.5 and 30.0 °C, Student’s *t*-test; 25.0, 32.5 and 35.0 °C, Welch’s *t*-test). Survival of nymphs was highest at 25.0 and 27.5 °C and decreased with increasing temperature, being lowest at 35.0 °C (Figure 4C).

**FIGURE 4.**
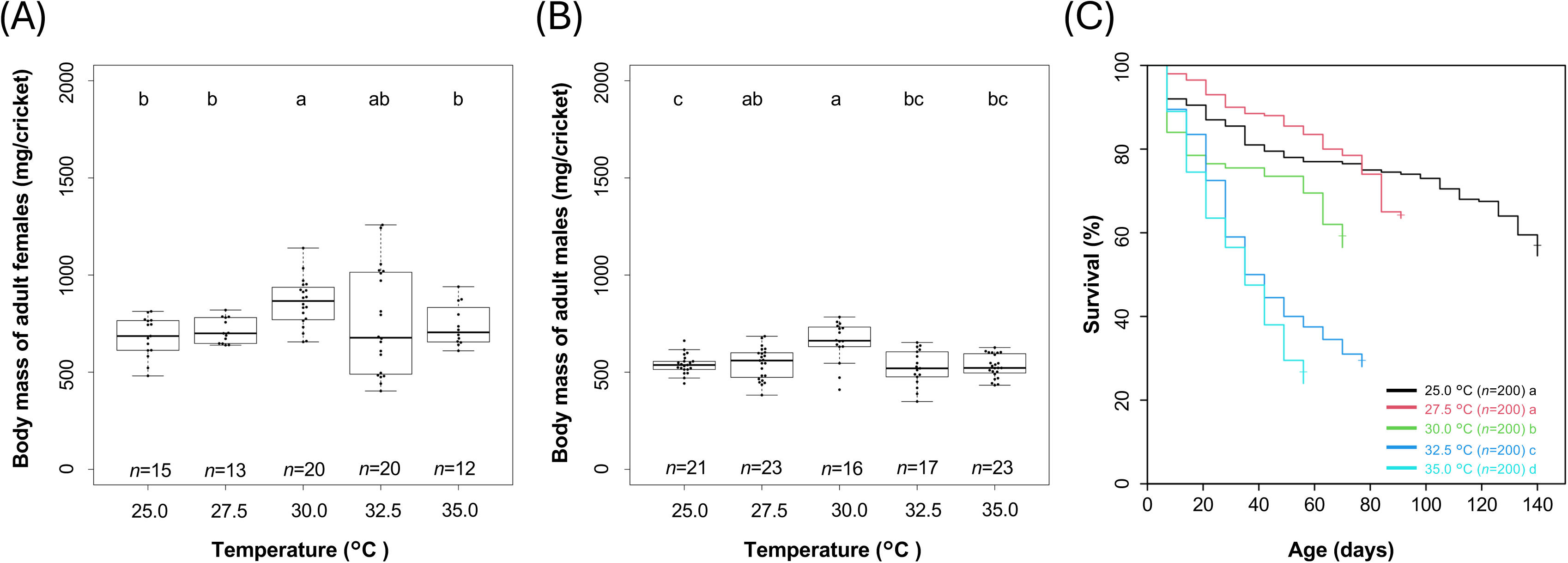
Growth performance of *Teleogryllus occipitalis*. (A) Female and (B) male adult body mass at each temperature (25.0, 27.5, 30.0, 32.5, and 35.0 °C) on the day of peak body mass (d133, d91, d70, d77, and d49, respectively). Different letters indicate significant differences among temperatures (*P* < 0.05). (C) Survival rate at each temperature from d0 to d140, the final recording day at 25.0 °C. Different letters indicate significant differences among survival curves (pairwise log-rank test, *P* < 0.05).

### Model simulation of PVH

In this study, the logistic model was used to calculate PVH. PVH differed among temperatures in both species (Figures 5 and 6). In *G. bimaculatus*, the highest PVH was predicted at 30.0 °C, reaching 13.82 kg/m^3^ at 38 days. In *T. occipitalis*, the highest PVH was also predicted at 30.0 °C, reaching 7.12 kg/m^3^ at 62 days. PVH values for all temperatures are listed in Table S2.

**FIGURE 5.**
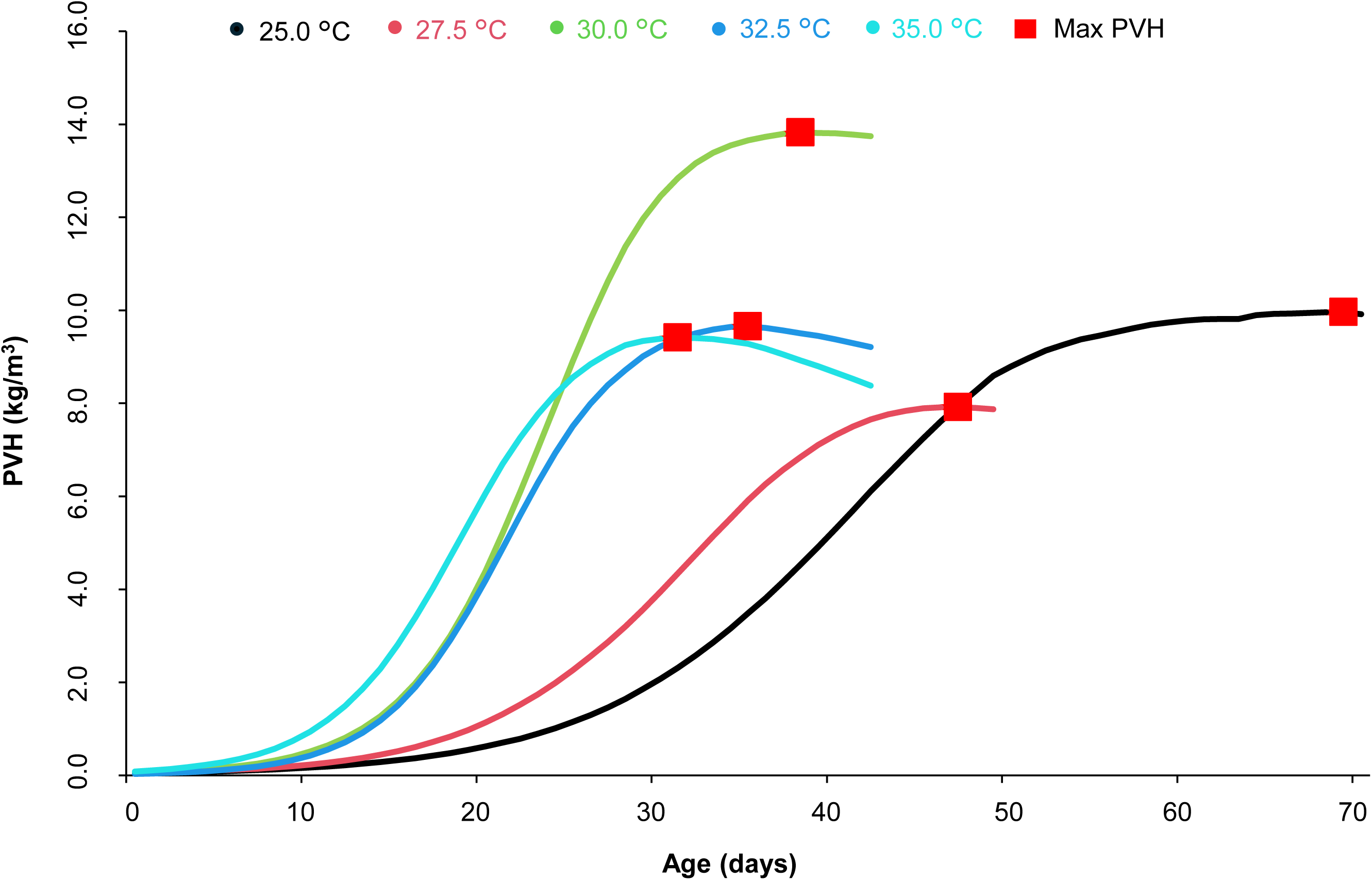
Predicted value of harvest (PVH) across temperatures in *Gryllus bimaculatus*. PVH was calculated by combining predicted body mass from the logistic growth model with observed survival probability linearly interpolated between weekly observations (Equation 4). Red symbols indicate the maximum PVH at each temperature.

**FIGURE 6.**
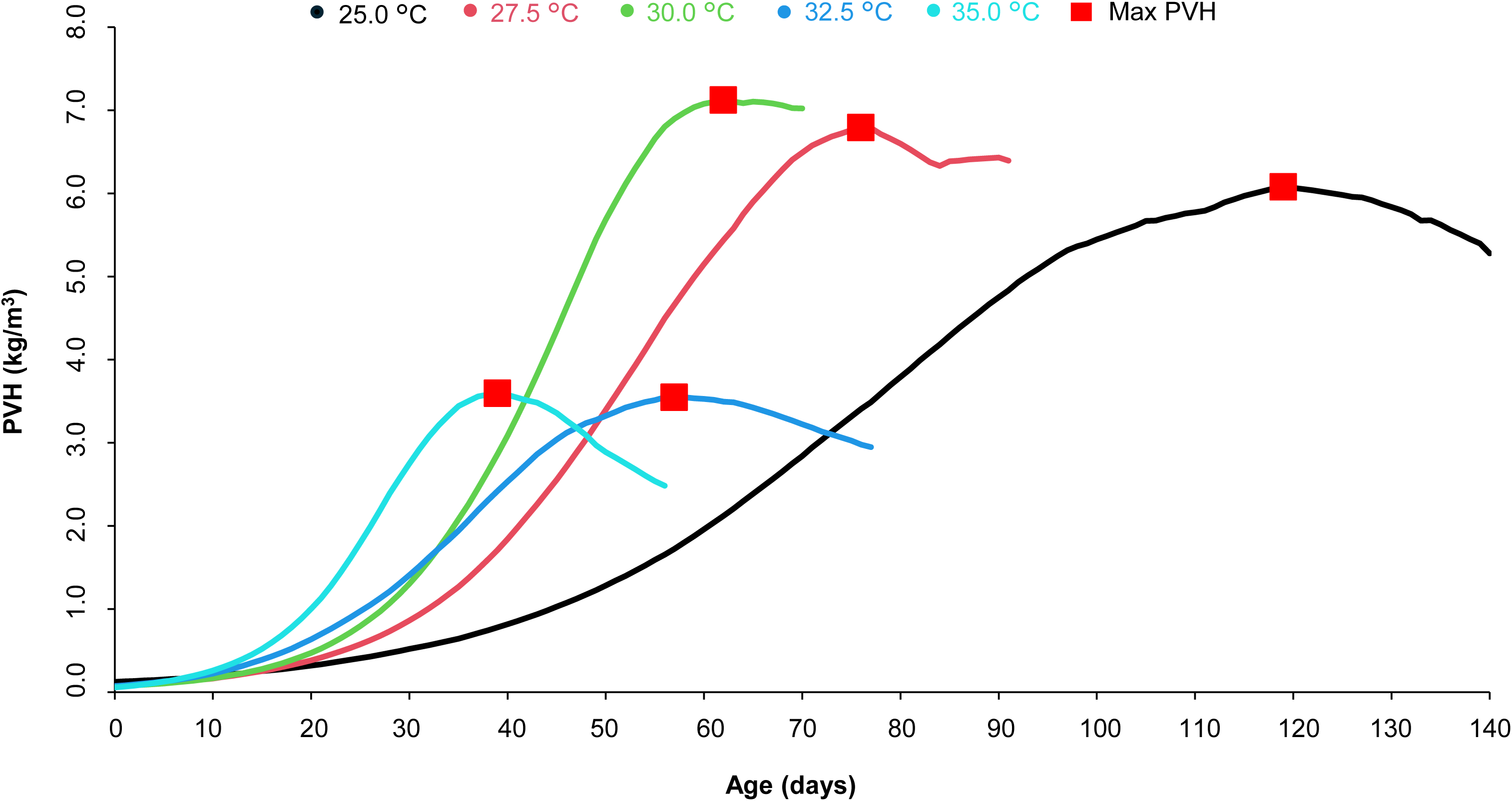
Predicted value of harvest (PVH) across temperatures in *Teleogryllus occipitalis*. PVH was calculated by combining predicted body mass from the logistic growth model with observed survival probability linearly interpolated between weekly observations (Equation 4). Red symbols indicate the maximum PVH at each temperature.

## Discussion

To date, studies of cricket growth have not compared multiple growth models. The three non-linear models used in this study fitted the data well at each temperature for both species, and the logistic and Gompertz models fitted particularly well. The differences in model-fitting accuracy across various ages and air temperatures are attributable to the distinct mathematical structures inherent in each growth model. Among the models we employed, the Gompertz is characterised by an inflection point occurring at approximately 36.8% of the asymptotic body mass; growth accelerates until this point and decelerates thereafter. In contrast, the logistic model defines its inflection point at exactly 50% of the asymptotic body mass, forming a symmetric growth curve in which the initial acceleration phase and the subsequent deceleration phase are balanced (Dhar and Bhattacharya, 2018; Goshu and Koya, 2013). The von Bertalanffy model has an earlier inflection point, occurring at approximately 29.6% of the asymptotic body mass. Thus, this model assumes that the maximum absolute growth rate is reached relatively early, followed by a comparatively long deceleration phase as body mass approaches its asymptotic value (Goshu and Koya, 2013).

As the rearing temperature has a strong impact on cricket growth, identifying the optimum conditions is important (Kong *et al*., 2025; Magara *et al*., 2024). The asymptotic body mass and growth rate of *G. bimaculatus* are temperature dependent, increasing from 20 to 32 °C and subsequently declining up to 37 °C (Magara *et al*., 2024). In this study, we analysed the growth characteristics at multiple temperatures using three non-linear growth models. The asymptotic body mass (*A*) peaked at 30.0 °C, whereas the growth rate constant (*k*) peaked at 35.0 °C except for the logistic model in *G. bimaculatus* (32.5 °C). These patterns are broadly consistent with the trend reported by Magara *et al*. (2024).

The reduction of adult body mass of *G. bimaculatus* at 25.0 °C compared with at 30.0 °C was observed both by Magara *et al*. (2024) and in our study (Figure 3) in both males and females even though there is a difference in the timing of weighing crickets (see below). The hypothesis that the asymptotic body mass is the same at 25.0 and 30.0 °C is rejected by our model estimates (Table 2, Figure 1). Therefore, the crickets reared at 30.0 °C grow not only faster but also larger than crickets at 25.0 °C. Body mass reduction at 35.0 °C compared with at 30.0 °C was observed both by Magara *et al*. (2024) and in our study in females (Figure 3A). Newly emerged female crickets (*A. domesticus*) are lighter than those 2 weeks after the final moult (Liao *et al*., 2025). This is due partly to the accumulation of fat body lipids (*Locusta migratoria*; Li *et al*., 2025) and ovarian development (*G. bimaculatus*; Lorenz, 2007) after the final moult. In other insect species, high temperature is reported to reduce the volume of fat bodies (*Drosophila melanogaster*; Klepsatel *et al*., 2016) and the ovary (*Anopheles gambiae*; Martin *et al*., 2025). Heat stress is therefore likely a factor in the body mass reduction in crickets reared at 35.0 °C. However, although we found that male crickets had a reduced body mass at 35.0 °C compared with at 30.0 °C, Magara *et al*. (2024) did not observe a significant reduction. The fact that newly emerged male crickets (*A. domesticus*) are lighter than crickets 2 weeks after the final moult (Liao *et al*., 2025) is due partly to the expansion of the testis (*Schistocerca gregaria*; Hiroyoshi *et al*., 2022) and the fat body (*Locusta migratoria*; Li *et al*., 2025) after the final moult. We agree with the hypothesis that heat stress suppresses expansion of the fat body (*D. melanogaster*; Klepsatel *et al*., 2016) and the testis (*Tribolium castaneum*; Sales *et al*., 2021). Magara *et al*. (2024) weighed their adults only 3 days after the final moult, probably before any negative effects of heat on the fat body and testes arose. In contrast, our adult crickets were more likely to have demonstrated heat shock effects (body mass reduction) because we weighed them from zero to several weeks after the final moult. Magara *et al*. (2024) reported that the body mass peak of female adults of *G. bimaculatus* was observed under a 32.0 °C treatment, whereas we observed the peak with a 30.0 °C treatment. The difference might have been derived from a difference in the population used for the experiments. A Kenyan population was used by Magara *et al*. (2024), whereas the initial origin of our population was unknown. In *Drosophila virilis*, variations in heat tolerance among populations of the same species have been observed (Yamamoto and Ohba, 1982). To date, data on sexual dimorphism in the body masses of adult crickets have been accumulated mainly for *A. domesticus*. Newly emerged *A. domesticus* females are heavier than males (Clifford and Woodring, 1990). In addition, females grow faster than males after the final moult owing to ovarian growth in adult female *A. domesticus* (Clifford and Woodring, 1990). In *G. bimaculatus*, the body mass of adult females 3 days after emergence was larger than that of males within a temperature range of 27.0 to 37.0 °C (Magara *et al*., 2024). Our results of *G. bimaculatus* and *T. occipitalis* do not contradict these previous studies. In our study, *T. occipitalis* showed a tendency in adult body mass similar to that of *G. bimaculatus*. Both species live in tropical to subtropical areas; *T. occipitalis* is distributed in South, Southeast and East Asia (He *et al*., 2017; Jaiswara *et al*., 2021) whereas a widely spread species *G. bimaculatus* is originally native to northern Africa and south-western Asia (Kulessa *et al*., 2025). Our results imply that both species are adapted to warm temperatures.

Survival was highest at 25.0 °C in both species, although it did not differ significantly from that at 30.0 °C in *G. bimaculatus* or from that at 27.5 °C in *T. occipitalis* (Figures 3C and 4C). Kong *et al*. (2025) found higher survival rates at lower temperatures in the tropical house cricket *Gryllodes sigillatus* reared at temperatures from 20.0 to 38.0 °C. Takacs *et al*. (2023) compared the survival of *A. domesticus* from day 20 to the first male chirping, a sign of adult maturity, among three rearing temperatures (25.0, 30.0 and 35.0 °C). They found that survival was significantly higher at 25.0 °C than at 30.0 °C, whereas survival at 35.0 °C did not differ from that at either temperature. Our results are partly consistent with the findings in these two species, although the temperature effect on survival was more pronounced in our study, particularly in *T. occipitalis*, possibly because of the longer rearing period from hatching. Under high ambient temperatures, insects, which are ectotherms, are exposed to the risk of stressors such as protein denaturation and oxidative stress, which can affect individual survival (Banfi *et al*., 2025). Insects then express heat shock proteins to minimise the effects of the heat stress (Banfi *et al*., 2025). Analyses of heat shock protein expression levels are required to reveal whether heat stress affects cricket mortality. Takacs *et al*. (2023) also quantified the abundance of *Acheta domesticus* densovirus but found no clear relationship between viral abundance and survival; viral abundance tended to be lower at 35.0 °C.

One potential application of this model is the prediction of productivity. To evaluate cricket productivity, we defined PVH as the predicted body mass of crickets per unit rearing volume at an arbitrary age, calculated from the predicted body mass and survival rate. The PVH analysis showed that 30.0 °C produced the highest maximum PVH in both species. In *G. bimaculatus*, the maximum PVH was 13.82 kg/m^3^ at 38 days, whereas in *T. occipitalis*, it was 7.12 kg/m^3^ at 62 days. These results suggest that 30.0 °C is the most suitable constant-temperature condition for maximising PVH in both species, although the optimal harvest timing differs between species. From the perspective of yield and time efficiency, *G. bimaculatus* appears to be more suitable for rapid production, as it reached a higher maximum PVH earlier than *T. occipitalis*. In contrast, *T. occipitalis* required a longer rearing period to reach its maximum PVH. In closed rearing systems at latitudes higher than the temperate zone, where winter occurs, maintaining high temperatures requires substantial energy input. Because temperatures higher than 30.0 °C did not increase the maximum PVH, there appears to be little benefit in setting the temperature above 30.0 °C throughout the production cycle. However, relatively high-temperature treatments increased PVH during the earlier stages of rearing. Therefore, higher temperature settings, such as 32.5–35.0 °C during the early stage followed by 30.0 °C rearing, may improve time efficiency, although heating costs must be considered. Further studies, including rearing experiments under variable temperature regimes throughout the crickets’ life cycle, will be required to confirm this hypothesis.

Crickets are produced worldwide, and in addition to the species targeted here, multiple others, such as *A. domesticus*, *T. mitratus*, and *G. locorojo*, are commercially reared, with variations in feed and rearing conditions (Magara *et al*., 2024; Ruang-Rit *et al*., 2025). Differences in rearing conditions result in variations in growth rate and final body mass (Sorjonen *et al*., 2019). By applying the modelling and calculation methods used here to each production region, rearing method and species, it will be possible to analyse growth characteristics.

## Conclusion

This study compared the growth characteristics of *G. bimaculatus* and *T. occipitalis* reared at five different constant temperatures using three non-linear growth models. The resulting growth predictions were integrated with survival probabilities to estimate yields at different temperatures and ages. The Gompertz, logistic, and von Bertalanffy models all fitted the data well. In both species, the asymptotic body mass was greatest at 30.0 °C and the growth rate constant was generally greatest at 35.0 °C. For both species, PVH was highest at 30.0 °C, reaching 13.82 kg/m^3^ at day 38 for *G. bimaculatus* and 7.12 kg/m^3^ at day 62 for *T. occipitalis*. These results indicate that 30.0 °C is the most suitable constant-temperature condition for maximising cricket productivity, although the optimal harvest age differs depending on the species. Analysis using non-linear growth models can not only compare growth characteristics but, when combined with observed survival rates, can identify the optimal rearing temperature and harvest timing. In the future, we will consider adapting this method to other environmental conditions.

## Supporting information

Supplemental Tabel S1

Supplemental Tabel S2

Supplemental Tabel S3

Supplemental Tabel S4

## Acknowledgements

We thank Dr Takashi Kuriwada of Kagoshima University for sharing the population of *T. occipitalis*.

## Data availability

The body mass and survival data supporting this study are provided in Tables S3 and S4. Other data are available from the corresponding author upon reasonable request.

## Supplementary materials

The following supplementary materials are available online. Figure S1 shows the rearing container used in this study, including the arrangement of the feed dish, watering device, paper towelling, egg carton shelters, non-woven fabric, and ventilated lid. Table S1 lists the ingredients and chemical composition of the experimental diet. Table S2 gives the initial parameter values used for non-linear least-squares fitting of each growth model and the daily PVH values at each temperature in both species. Tables S3 and S4 contain the body mass and survival data recorded for *G. bimaculatus* and *T. occipitalis*, respectively.

