## Supplementary figures and images for "Non-linear growth models combined with survival rates identify the optimal rearing temperature and harvest timing in edible crickets, *Gryllus bimaculatus* and *Teleogryllus occipitalis*"

### Supplemental Figure S1

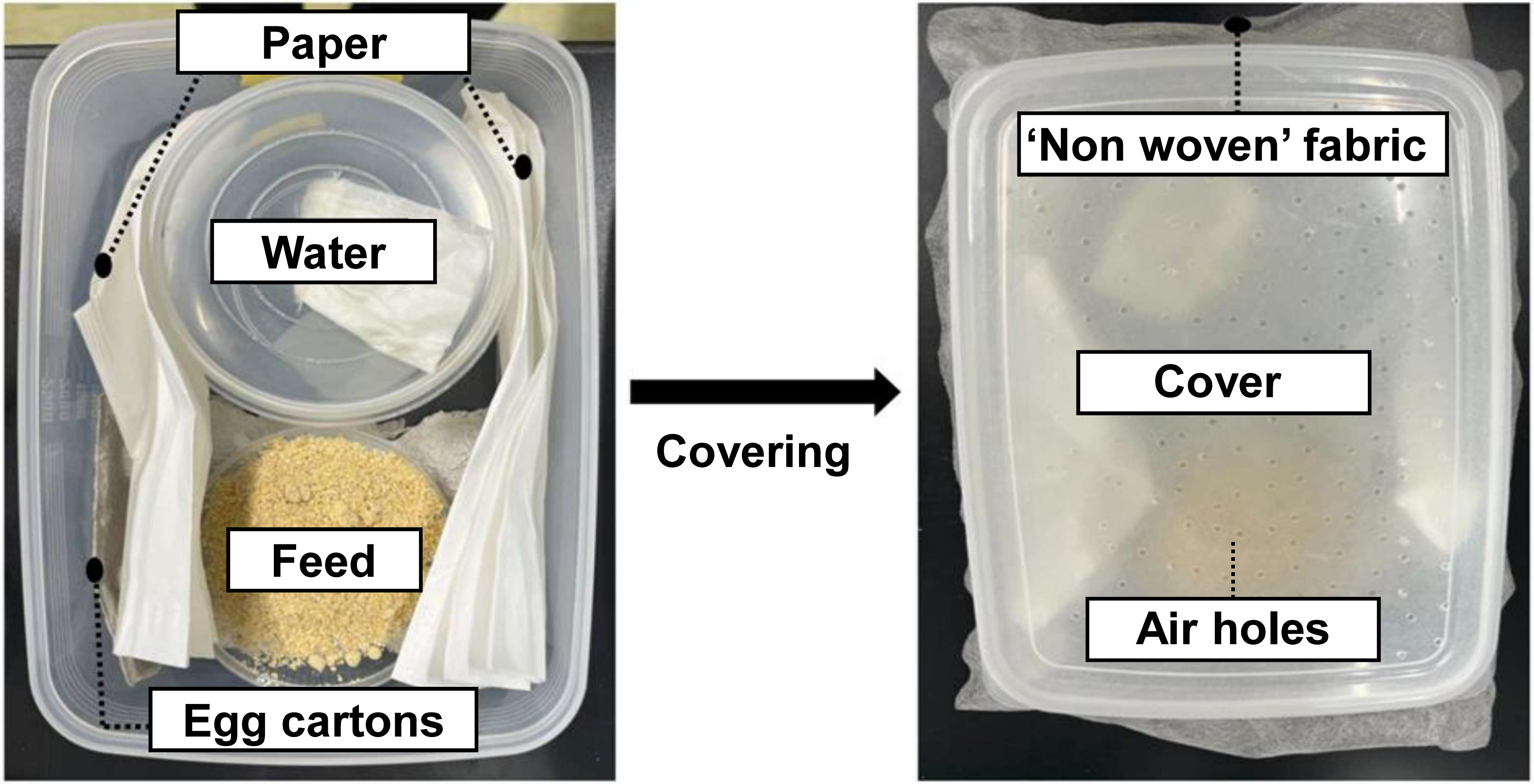
