## Supplemental Tabel S1 for "Non-linear growth models combined with survival rates identify the optimal rearing temperature and harvest timing in edible crickets, *Gryllus bimaculatus* and *Teleogryllus occipitalis*"

**Supplementary material**

**TABLE S1 Feed ingredients and chemical composition of the experimental diet; means ± SD.**

| Ingredient (%) | |
| --- | --- |
| Corn (maize) ^1^ | 45.50 |
| Soybean meal ^2^ | 21.00 |
| Rice bran ^3^ | 14.00 |
| Full-fat soybean ^4^ | 10.00 |
| Fishmeal ^5^ | 5.00 |
| Calcium carbonate ^6^ | 2.00 |
| Soybean oil ^7^ | 1.00 |
| Calcium phosphate, monobasic ^8^ | 0.50 |
| Sodium chloride ^9^ | 0.50 |
| Premix ^10^ | 0.50 |
| Total | 100.00 |
| Nutrient composition (%) ^a^ | |
| Moisture | 10.45 ± 0.23 |
| Protein | 21.96 ± 0.18 |
| Fat | 8.44 ± 0.10 |
| Carbohydrate | 51.70 ± 0.42 |
| Ash | 7.46 ± 0.25 |

^1^(kotubu-5kg; Saitou Corporation, Gunma, Japan), ^2^(TIS-DK002r; Tamagoya, Ibaraki, Japan), ^3^(B0F6JZRHBY; SI system, Japan), ^4^(Zesshi daizu; Sankosha, Fukushima, Japan), ^5^(TKE-FSM60002; Tamagoya, Ibaraki, Japan), ^6^(030-00385; FUJIFILM Wako Pure Chemical Corporation, Osaka, Japan), ^7^(B0076JR7C2; Rikennosankakou, Fukuoka, Japan), ^8^ (Yoneyama Chemical Industry, Osaka, Japan), ^9^(191-01665; FUJIFILM Wako Pure Chemical Corporation, Osaka, Japan)^, 10^Vitamin and mineral premix per kg of diet (Kumiai nyuu VM; JA zen-noh kumiaisiryou, Tokyo, Japan): vitamin A, 3,000,000 IU; vitamin D3, 600,000 IU; vitamin E, 3.00 g; vitamin K3, 0.50 g; vitamin B1, 0.40 g; vitamin B2, 3.00 g; vitamin B6, 1.00 g; vitamin B12, 2.50 mg; Nicotinamide, 5.00 g; folic acid, 0.125 g; calcium-D-pantothenate, 2.00 g; Choline chloride, 120.00 g; FeSO4, 25.21 g; CuSO4, 2.39 g; CoSO4, 41.79 mg; Mn, 24.65 g; Zn, 24.51 g.

^a^The nutrient composition of the feed was analysed by using the following standard methods: Protein was determined by the macro Kjeldahl method, fat by the Soxhlet extraction method, carbohydrate by the difference method, ash by the direct ashing method, and moisture by the oven drying method at ambient temperature.
